# Evolutionary responses to conditionality in species interactions across environmental gradients

**DOI:** 10.1101/031195

**Authors:** Anna M. O’Brien, Ruairidh J.H. Sawers, Jeffrey Ross-Ibarra, Sharon Y. Strauss

**Affiliations:** Dept. of Ecology and Evolutionary Biology, University of Toronto, Toronto, ONT, Canada M5S 3B2; Center for Population Biology, University of California, Davis, CA 95616; Dept. of Plant Sciences, University of California, Davis, CA 95616; Dept. of Evolution and Ecology, University of California, Davis, CA 95616; Laboratorio Nacional de Genómica para la Biodiversidad (LANGEBIO), Centro de Investigación y de Estudios Avanzados del Instituto Politécnico Nacional (CINVESTAV-IPN), Irapuato, 36821, Guanajuato, Mexico; Genome Center, University of California, Davis, CA 95616

**Keywords:** biotic interactions, mutualism, local adaptation, co-adaptation, environmental gradients

## Abstract

The outcomes of many species interactions are conditional on the environments in which they occur. A common pattern is that outcomes grade from being more positive under stressful conditions to more antagonistic or neutral under benign conditions. The evolutionary implications of conditionality in interactions have received much less attention than the documentation of conditionality itself, with a few notable exceptions. Here, we predict patterns of adaptation and co-adaptation between partners along abiotic gradients, positing that when interactions become more positive in stressful environments, fitness outcomes for mutations affecting interactions align across partners and selection should favor greater mutualistic adaptation and co-adaptation between interacting species. As a corollary, in benign environments, if interactions are strongly antagonistic, we predict antagonistic co-adaptation resulting in Red Queen or arms-race dynamics, or reduction of antagonism through character displacement and niche partitioning. We predict no adaptation if interactions are more neutral. We call this the CoCoA hypothesis: (**Co**)-adaptation and **Co**nditionality across **A**biotic gradients. We describe experimental designs and statistical models that allow testing predictions of CoCoA, with a focus on positive interactions. While only one study has included all the elements to test CoCoA, we briefly review the literature and summarize study findings relevant to CoCoA and highlight opportunities to test CoCoA further.

## Outcomes of biotic interactions depend on abiotic conditions

The fitness impacts of biotic interactions are shaped by the conditions in which they occur. For example, warming temperatures cause corals to expel their zooxanthellae symbionts (Hoegh-Guldberg, 1999), increasing fire frequency and severity favors invasive over native grasses in competitive interactions (D’Antonio and Vitousek, 1992), and predation on pepper moths is altered by the prevalence of air pollution (Kettlewell, 1955). Conditionality in mutualisms is also well known (Cushman and Whitham, 1989; Bronstein, 1994), and a meta-analysis of mutualism studies finds that mutualistic outcomes are variable across space and habitats (Chamberlain et al., 2014).

Two well-developed and related models of species interactions seek to predict changing fitness impacts of biotic interactions for partners (interaction outcomes) across gradients. First, economic models of mutualism describe inequalities with respect to resources and predict conditional outcomes from true mutualistic outcomes (both species receive fitness benefits, or +,+ outcomes) to antagonism (+,- or -,- fitness outcomes). When the resources a participant *receives* in trade from partners are those that are most limiting to the participant’s fitness, the benefits from trading are maximized; when resources the participant *provides* to partners limit the participant’s fitness, the costs of engaging in trade are maximized (Johnson, 1993; Schwartz and Hoeksema, 1998; Bever, 2015). Resource-based conditionality has been shown to exist for many “mutualisms” (Bronstein, 1994), for example between plants and mycorrhizal fungi, which typically provide soil nutrients to plants in exchange for carbon. This exchange benefits plants in low nutrient (stressful) conditions, but often imposes costs when nutrient availability is high (Smith et al., 2010).

A second model closely tied to environmentally conditional outcomes in species interactions is the Stress-Gradient hypothesis (SGH). The SGH posits that the relative importance of costs and benefits from biotic interactions changes across stress gradients (Bertness and Callaway, 1994), and that interactions will gradually shift from having neutral or negative outcomes under benign abiotic conditions to having beneficial outcomes under stressful conditions (Brooker and Callaghan, 1998; Malkinson and Tielbörger, 2010). In some cases, plants are mutualistic as seedlings in stressful conditions, but are less affected by these stresses as adults, and they then compete (Sthultz et al., 2007). In other cases, stresses may be sufficiently great that the positive interactions between species are maintained through the lifecycle. For example, stressful high altitude conditions often result positive interactions between species that are positive throughout life (Sthultz et al., 2007; He et al., 2013). For the purposes of this paper, we consider cases in which the conditionality of abiotic stress is either consistent over the lifetime of an interaction (e.g., seedling to adult), or we simplify to the net fitness effects of the interaction. In other words, if seedlings of different species facilitate each other, and seedling mortality has the greatest effects on fitness, then, even if adults compete, we would consider the interaction under stress as positive. A meta-analysis of SGH in plants found consistent shifts towards facilitation (0,+) or reduced competition (0,- or -,- with smaller fitness effects) at high stress (He et al., 2013).

These separate theories are united by a focus on change in interaction benefits over abiotic gradients: when interactions ameliorate fitness-limiting factors, they are expected to have positive effects on fitness, and when they exacerbate fitness-limiting factors, they should decrease fitness. The theories use different language for overlapping concepts (“stress”,“limiting resources”). Here, we use “stress” to describe this overlap: an abiotic condition that limits fitness. The SGH and resource-based conditionality were originally detailed to explain changes from competition to facilitation in plant interactions and changes from mutualism to antagonism in plant-microbe interactions, respectively, yet they have been applied to a diversity of interactions such as detritivore-detritivore (Fugere et al., 2012), herbivore-herbivore (Dangles et al., 2013), plant-herbivore (Daleo and Iribarne, 2009), and bacterial cross-feeding (Hoek et al., 2016), all of which become increasingly facilitative or decreasingly costly as a stress the interaction ameliorates increases.

The evolutionary implications of conditionality in interactions have received much less attention than the documentation of conditionality itself, with notable exceptions (Schwartz and Hoek-sema, 1998; Thompson, 2005; Bronstein, 2009; Michalet et al., 2011). The geographic mosaic theory of coevolution (GMTC,Thompson, 2005) suggests that as fitness consequences of interactions vary across space, selection pressure from these variable interactions will result in different evolutionary outcomes. The GMTC is well supported (Thompson, 2005; Schemske et al., 2009), yet lacks a framework for linking characteristics of the environment to specific evolutionary outcomes.

Here, we generalize these predictive frameworks for species interaction outcomes and unite them with evolutionary principles. Our hypothesis links effects of limiting gradients on interaction outcomes to the degree of adaptation to species interactions in pairs of populations across stress gradients. We first leverage existing theory of conditionality, stress gradients, and geographic mosaics to generate predictions, then propose experimental and analytical methods for testing this hypothesis and discuss existing relevant literature.

## Evolutionary responses to conditionality: a hypothesis

Because conditionality models predict that environmental or resource gradients result in predictable variation in interaction outcomes, we suggest that evolution in these contexts might also result in predictable outcomes. Extending the predictions of conditionality in interaction outcomes to coevolutionary dynamics, we predict selection should result in adaptation and coadaptation in species interactions that are shaped by environmental gradients.

Where the interaction ameliorates a fitness limiting stress in both species, mutations in one species that reduce stress on a partner species can simultaneously increase fitness in both species. The increase in fitness of the partner species increases the frequency or extent of the interaction for the first species, ameliorating more stress and positively impacting fitness. This phenomenon is known as fitness feedback (Sachs et al., 2004), and such mutations will be favored by selection. Genetic variation in the traits of one partner that ameliorate stress in the other should thus impact fitness of both partners in these stressful sites. As selection continues to fix mutations ameliorating the stress of partners, we predict increasing mutual benefit at stressful or resource-limited ends of environmental gradients due to fixation of mutations in both partners (mutualistic co-adaptation) or just one partner alone (mutualistic adaptation, Figure 1).

**Figure 1:**
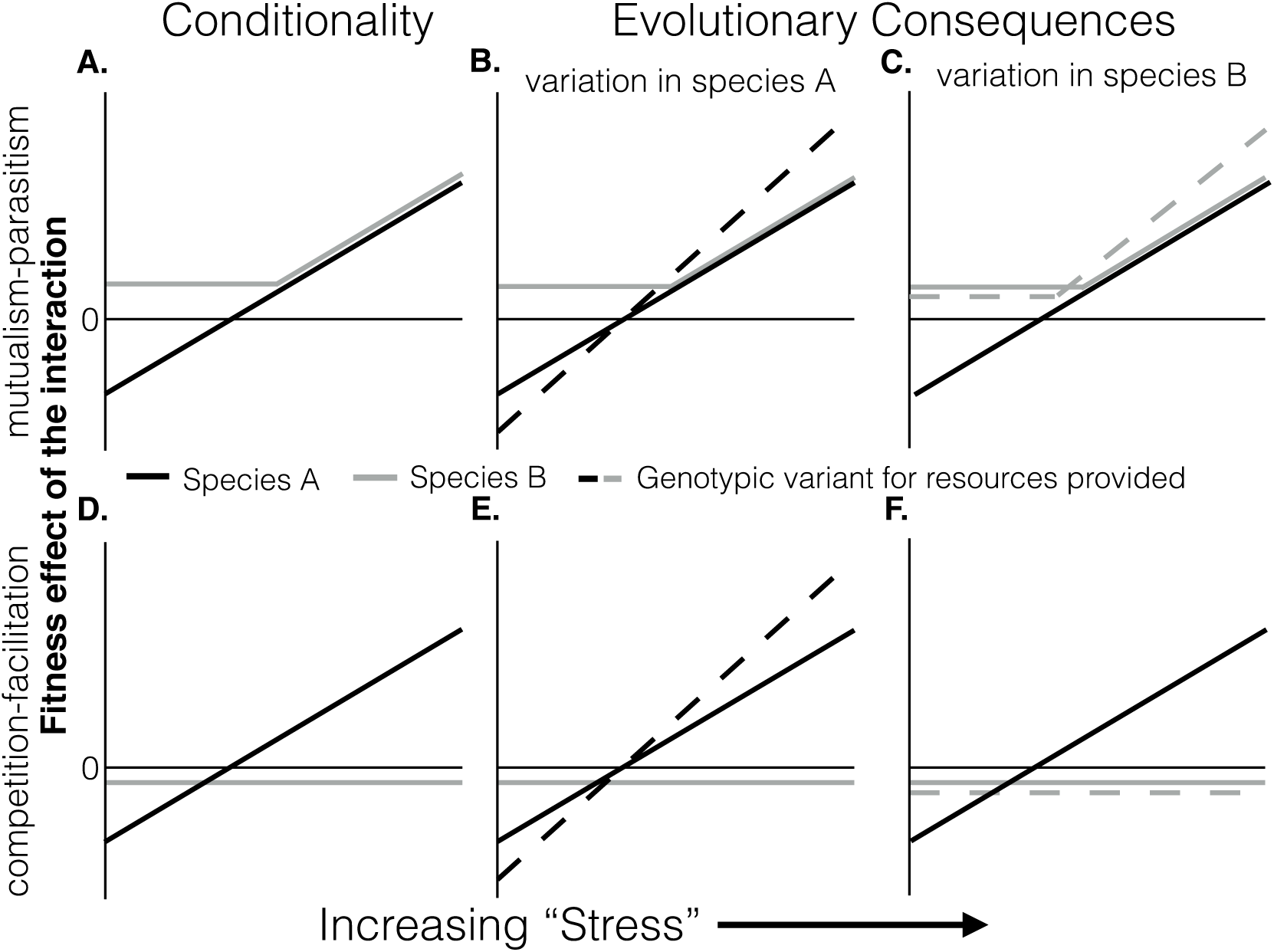
Conditionality hypotheses (A,D) and CoCoA predictions (B,C,E,F) at low and high stress. A-C, from parasitism to mutualism; D-F, from competition to facilitation. A & D, interaction outcomes shift towards more positive interaction outcomes at more stress-limited sites, as predicted by generalizing either SGH or limiting resource conditionality. B, E, the fitness of a variant of species A (the species parasitized at low stress in B and facilitated in E) that provides more benefit to species B (the species parasitic at low stress in B and the facilitator in E) across interaction types and shifts is now depicted next to the original genotype in dashed line. C,F, the same, but for variation in species B. Depending on the interaction type and stress, the selection on species A and B would favor the variant, original genotype, or neither, but variants are more favored (or less disfavored) at higher stress. CoCoA thus suggests increasing mutualistic local co-adaptation or adaptation at high stress sites, and where interactions grade into increasing antagonism (+,- or -,-), increasing antagonistic co-adaptation (for +,-) or adaptation to avoid interactions (for -,-) is favored.

At the ends of gradients that are “benign” with respect to stresses or resources, fitness will be instead limited by either costs of the interaction or by unrelated factors. Interactions between species may become neutral or shift towards antagonism (Johnson, 1993; Bertness and Callaway, 1994; Schwartz and Hoeksema, 1998), which we predict will result in a variety of coevolutionary outcomes.

If the interaction is neutral for one or more partners, we predict no co-adaptation, though if the interaction continues to negatively or positively impact fitness of one partner, adaptation in this partner will still be influenced by the interaction. For example, in shifts of plant-plant competition towards facilitation with increasing stress, facilitation is often not mutual (He et al., 2013; Schob et al., 2014b,a). When interactions do not alter fitness, mutations that increase investment in interactions will drift, or will be removed by selection if the investment itself is costly to produce.

When the interaction is antagonistic in benign conditions (-,- or +,-), the interaction may again strongly affect fitness, now inflicting high costs on one or both partners (Figure 1). Reciprocal selection in mutually antagonistic interactions (-,- as in many competitive interactions) could act either to reduce antagonistic interactions through avoidance of the interaction entirely (such as character displacement, Pfennig and Pfennig, 2009), or to avoid fitness costs through tolerance (Bronstein, 2009). Both of these responses to antagonistic interactions reduce the effects of the interaction on fitness, and reduce the strength of selection imposed by each species. Asymmetric antagonisms (+,-), such as trophic interactions (e.g. parasitism, predation), can result in asynchronous or oscillating Red-Queen coevolutionary dynamics such as arms-races (Toju et al., 2011) or frequency-dependent selection (Decaestecker et al., 2007). In particular for arms-races, this intensified coevolution in benign conditions will escalate offensive and defensive traits to more extreme values (Hochberg and van Baalen, 1998; Benkman et al., 2003; Hanifin et al., 2008). Mutations affecting asymmetric interaction outcomes will have high fitness consequences and will either swiftly fix or could exhibit cyclical dynamics under frequency-dependent selection.

Evidence exists that many traits affecting interaction outcomes have a genetic basis and can respond to selection. For example, variation in mutualistic benefit provided has been shown to have a genetic basis in many systems (e.g. Moran, 2001; Eaton et al., 2015; Klinger et al., 2016; Batstone et al., 2017) as has variation in resistance to antagonists (e.g. Staskawicz et al., 1995; Lively and Dybdahl, 2000; Decaestecker et al., 2007), and thus both can be expected to respond to selection.

Both theoretical and empirical work suggest that as the strength of selection on beneficial or antagonistic interactions increases, mutations improving interaction outcomes with local partners are more likely to fix (Parker, 1999; Nuismer et al., 2000; Kawecki and Ebert, 2004; Thompson, 2005; Schemske et al., 2009). Strong selection coupled with low gene flow is predicted to result in specific adaptation or co-adaptation between local populations. While extremely high gene flow would prevent adaptation along any gradient, intermediate gene flow could preclude local adaptation/co-adaptation within populations and instead promote general adaptation/coadaptation among sets of populations. We define specific benefit mutations as those that are specific to the genotypes of local partners (“specific benefits”). Specific-benefit mutations should fix under low gene flow while mutations underlying benefits to and from multiple partners (“generalized benefits”) are predicted to be favored when gene flow between stressful sites is higher (see also, “Interpretation of Results”, Figure 2).

**Figure 2:**
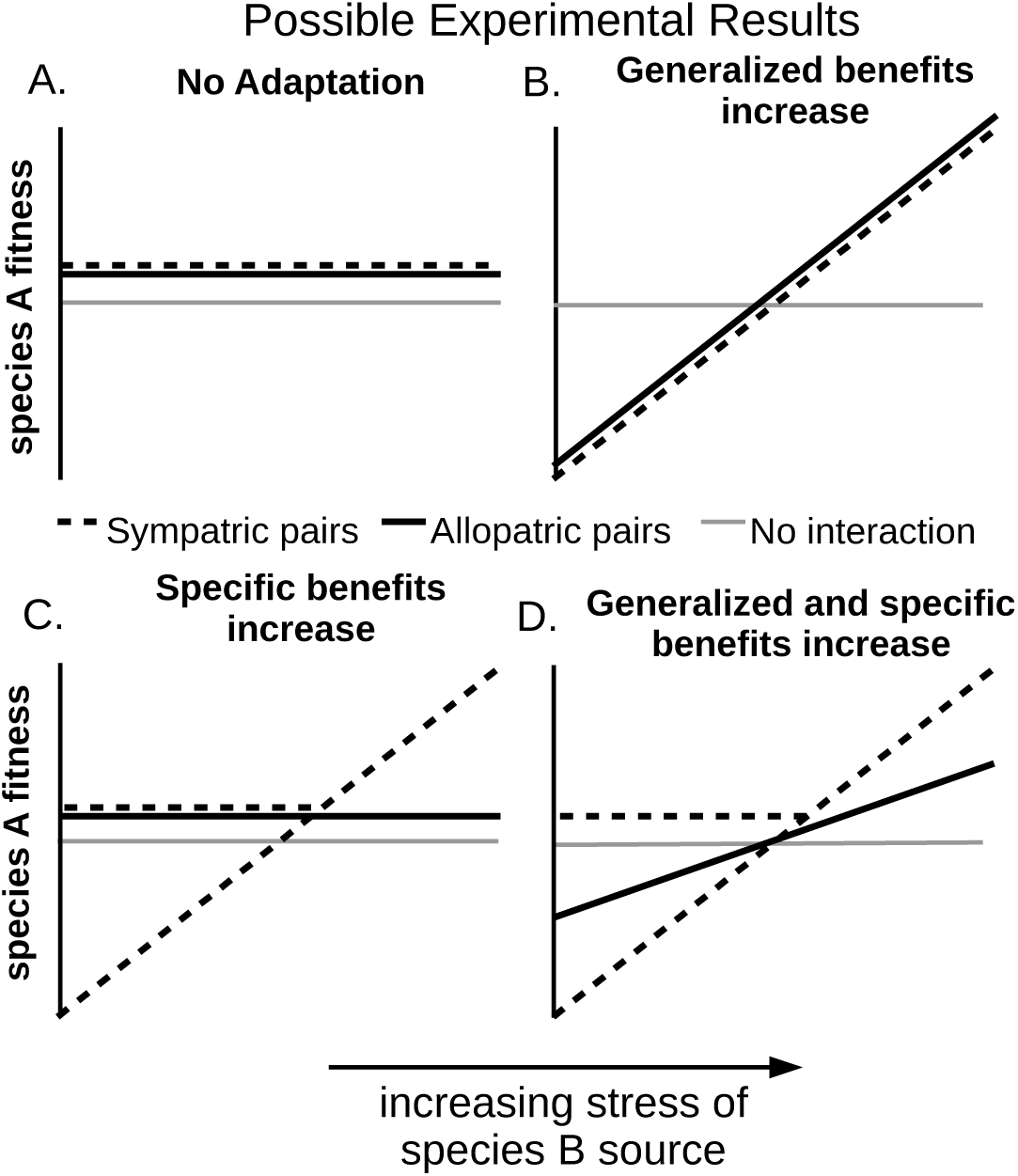
Possible experimental outcomes under high stress conditions. CoCoA predicts greater generalized fitness benefits provided by partners sourced from stressful sites across allopatric (solid lines) and sympatric (dashed lines) combinations (B, D). CoCoA predicts increasing specific fitness benefits of sympatric combinations with source stress (increasing difference of dashed and solid lines, C, D). For combinations with partners from benign sites, CoCoA predicts variable outcomes (multiple dashed lines), and no (A,C,D) or negative (B-D) sympatric effects. Increasing costs of sympatric partners as environments become more benign would indicate antagonistic adaptation, but might only be observed for one species, and other possibilities exist (see text). With antagonistic co-adaptation, a more likely result is high variance among population sympatric effects, if population pairs are at different stages in Red-queen dynamics. However, high variance in sympatric effects could alternately result from drift, and so is not a useful test. Without co-adaptation or adaptation, CoCoA expects no sympatric effects or difference across gradients (A). As a reference point, species A fitness without interacting with species B is shown in grey.

In sum, we predict that interactions with net fitness effects that shift in sign or strength along gradients will generate the most adaptation or co-adaptation near gradient extremes and least midrange, where neutral or reduced fitness impacts on one or more species prevent feedbacks. We predict evolution towards increasing mutualism and/or greater mutualistic co-adaptation where the interaction most ameliorates fitness-limiting stress. In contrast, we predict antagonistic evolutionary dynamics at benign sites, where interaction outcomes are expected to be more antagonistic. We call this the (**Co**)-adaptation to **Co**nditionality across **A**biotic gradients hypothesis, or CoCoA.

Below, we discuss designs that can test CoCoA. In designing a test for CoCoA, we focus primarily on interactions that are mutualistic either along part or the full length of the stress gradient, as we predict the coevolutionary outcomes will be consistent over time in mutualistic zones of the gradient, making these populations most straightforward to test at a single timepoint. In contrast, antagonistic coevolution is predicted in more benign conditions. It is well-known that antagonistic coevolution is difficult to test, as many patterns are consistent with, but not indicative of, antagonistic coevolution (see e.g. Lively and Dybdahl, 2000; Nuismer et al., 2000; Nuismer, 2006; Gandon et al., 2008; Frederickson, 2013; Stuart and Losos, 2013).

## Testing for CoCoA

Tests of CoCoA should include: (1) evidence of an environmental gradient that ranges from limiting to non-limiting for one (only adaptation predictions for the limited species are relevant) or both (all CoCoA predictions) partners; (2) evidence that the net fitness impact of the interaction on partners changes across the gradient due to changes in stress limitation; (3) measures of fitness outcomes in interactions with local and non-local partner pairs sourced from populations across the gradient. Throughout, we refer to populations of each species from the same site as sympatric and populations from different sites as allopatric. Measurements of partner effects on fitness must include both sympatric and allopatric partners to test for both generalized and specific benefits. Specific benefits (see above) could arise from specific populations of both species co-adapting to the other (specific co-adaptation), or only one species population adapting to the other if the interaction is +,0 (specific adaptation). Generalized benefits would arise if heritable traits adaptively increased the benefits provided to any partner (i.e. across multiple populations) at stressful sites (generalized adaptation or co-adaptation). Below we outline experimental designs and models to test CoCoA, and discuss interpretations of results.

### Experimental design

The ideal test of CoCoA will quantify two things. First, it will quantify the effects of the interaction on the fitness of both species sampled from across the gradient. Despite widespread documentation of conditionality (Chamberlain et al., 2014), abiotic predictors of conditionality remain unclear for many species interactions (for example, Maron et al., 2014). Second, the ideal test will quantify the extent of generalized and specific benefits between partner species across the gradient. For illustration, we provide an example test for the interaction between two species (Species “A” and “B”) along a gradient from stressful conditions, where CoCoA and conditionality hypotheses predict that species will mutually enhance each others’ fitness, to conditions where at least one species is predicted to have a negative effect on the other. CoCoA applies to other interactions that may span different outcome ranges across limiting stresses (e.g. competitive in benign sites to commensal in stressful sites), which can be tested in the same fashion. Except for interactions that never become commensal or mutually positive, the single timepoint tests are sufficient.

Testing CoCoA requires sampling populations of both species at sites along an identified stress gradient using the general approach proposed by Blanquart et al. (2013). To this approach, we add sources of populations across a gradient, and inclusion of gradient effects on fitness outcomes in the analyses. More populations always improves power, since population source site is the experimental unit, yet the number of populations must be balanced with the replication needed for each comparison. Under CoCoA, we predict increased generalized and specific benefits accruing from adaptation of partners at the stressful end of the gradient. In order to test for generalized adaptation (Figure 2, B and D, solid lines), one can regress the effect of Species B source population on Species A fitness across all populations of Species A sampled along the gradient. For example, a significant positive global effect on Species A fitness from interacting with a single population of Species B indicates that selection has favored generalized mutualistic traits in that Species B population. To quantify specific adaptation or co-adaptation between local populations of partners, it is necessary to assess the relative benefits received by both Species A and Species B with sympatric partners versus allopatric partners across the gradient (Figure 2, C and D, difference between dashed and solid lines).

While these comparisons may be made using all possible combinations of interacting partner populations of Species A and B, a fully crossed design is not required. We suggest designs that have twice as many allopatric as sympatric comparisons across the gradient. Power to estimate local adaptation in sympatry is maximized when the number of allopatric and sympatric comparisons are equal, and with the largest feasible number of populations (Blanquart et al., 2013). However, because our model includes a formal gradient term, and interactions with that gradient, our design requires additional allopatric comparisons relative to the number of sympatric comparisons. Paired populations of both Species A and B must be sampled from the same sites, and should be sampled to span the gradient, including intermediate sites, as stress is modeled as a continuous gradient in our approach. Sampling of paired populations across the gradient allows allopatric comparisons for each population from a source site of similar stress level, which increases the power to estimate change in sympatric effect across source site stress.

Random experimental combination of sampled populations will increase several biases in estimating allopatric effects. Populations sourced from the lowest stress sites will be more often combined with partners from sites with higher stress (intermediate and high) than other low stress sites (and vice versa). Populations from the highest and lowest stress sites will have a larger range in the difference in stress between their own source site and comparison population sites. A variety of designs minimize potential biases, and we provide one example in Figure 3.

**Figure 3:**
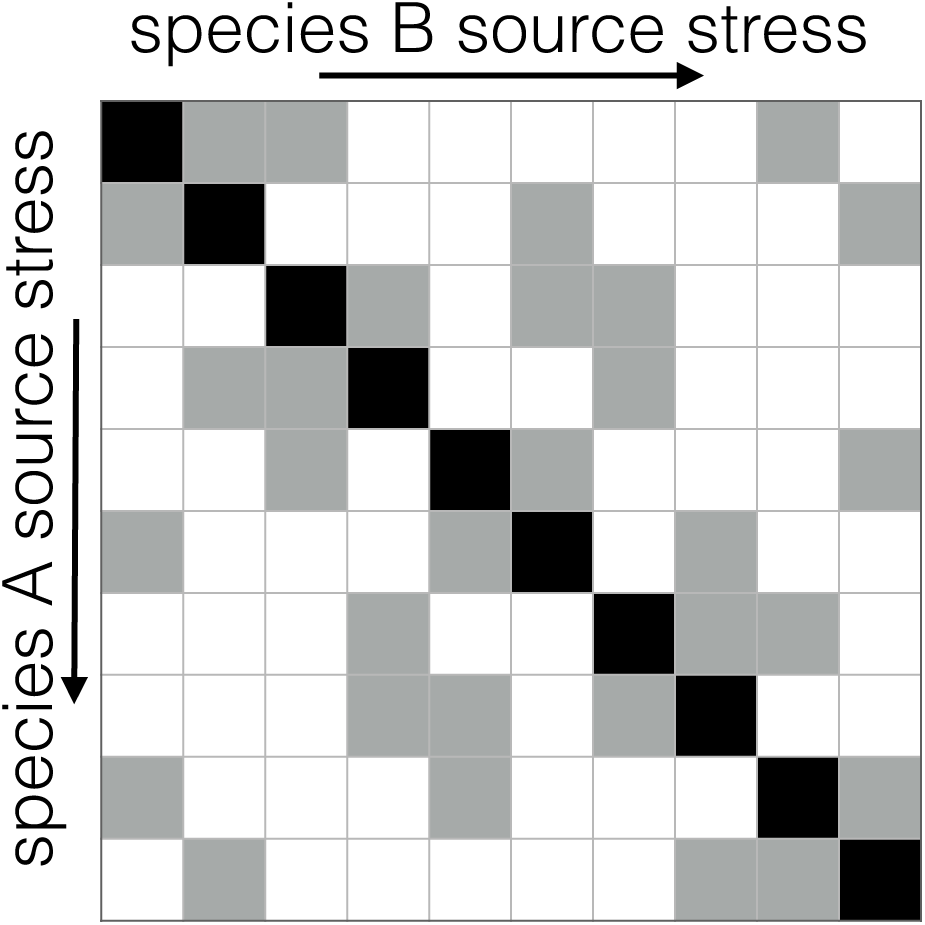
Possible sampling design and experimental combinations. Species A sources are in rows, arranged by increasing stress of source site from top to bottom. Species B sources are in columns, arranged by increasing stress of source site from left to right. Filled in squares are experimentally paired populations of A and B, including twice as many allopatric (grey) as sympatric (black) comparisons, and spreading sympatric and allopatric comparisons along the stress gradient for sources of both species A and species B.

Experiments should be run under conditions representative of those observed in natural populations, as inappropriate conditions may alter expressed benefits or costs of associating with partners (Lau and Lennon, 2012). Ideally, fitness measures will be as close to absolute fitness as possible, such as number of viable offspring. Running the experiment in multiple common gardens with different conditions allows a test of the CoCoA prerequisite that increasing stress shifts interaction outcomes towards increasing mutualism at high stress. While repeating in multiple environments is optimal, it may be possible only in systems where massive replication is feasible, such as in microbe-microbe interactions. Field environments have the benefit of being a more realistic context in which to test for co-adaptation between local populations, but are often constrained in replication both for the number population sources and the number of common garden (sensu lato) sites. We propose that a minimal design tests outcomes under conditions representing the stressful end of the gradient, which should maximize detection of mutualistic adaptation or co-adaptation. Here, our experimental design and analysis tests CoCoA predictions for this stressful region of the gradient (e.g., under reduced resources, water availability, etc.).

### A linear model framework

In classic tests of local adaptation, populations and sites are treated as discrete entities (Kawecki and Ebert, 2004; Blanquart et al., 2013). Our approach to detect local adaptation along a gradient uses a continuous approach to analyze gradient effects. We suggest modeling effects of partners and environments on fitness in a linear framework (non-linearity discussed below), where fitness in one focal partner at a time is the response variable *Y* (below), and then repeating across the other partner so that species A and B fitnesses are response variables in separate models. This linear testing framework defines generally better and worse mutualists using average fitness benefits conferred to partners across partner combinations, which follows recent advances in theory (Frederickson, 2013; Jones et al., 2015). Below we show Species A fitness as the response (*Y_A_*); the model for Species B fitness would be specified by swapping all A and B terms.

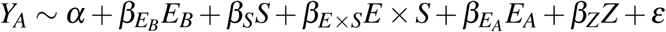

The estimated parameter for the main effect of source environment of the non-focal partner (here, the environment of Species B population source, *E_B_*, parameter *β*_EB_) is a test of the CoCoA prediction that Species B sourced from more stressful sites might be generally more mutualistic for all Species A populations than Species B sourced from the less stressful parts of the gradient. CoCoA predicts that *β_E_B__* should be positive.

Models include a slope parameter for the binary term (*S*) indicating whether origins of the interactors are sympatric (*S* = 1) or allopatric (*S* = 0) in addition to the slope parameter for the interaction between sympatry and the environmental gradient of source (*β_E×S_*), because we predict sympatry effects to vary across the gradient. Parameter estimates for effects of non-focal partner source environments (*β_E_B__*) compared to estimates for the environment interaction with sympatry (an environment × sympatry interaction denoted as *E × S*) allow us to tease apart generalized benefits from specific benefits along the gradient (Figures 2 & 4). CoCoA predicts that without extensive gene flow between high stress sites, *β_E×S_* should be positive; specifically that benefits accrued by sympatric partners from most stressful sites should be relatively greater than the benefits accrued by sympatric partners from other parts of the gradient, e.g. specific benefits are increased for stressful sites.

**Figure 4:**
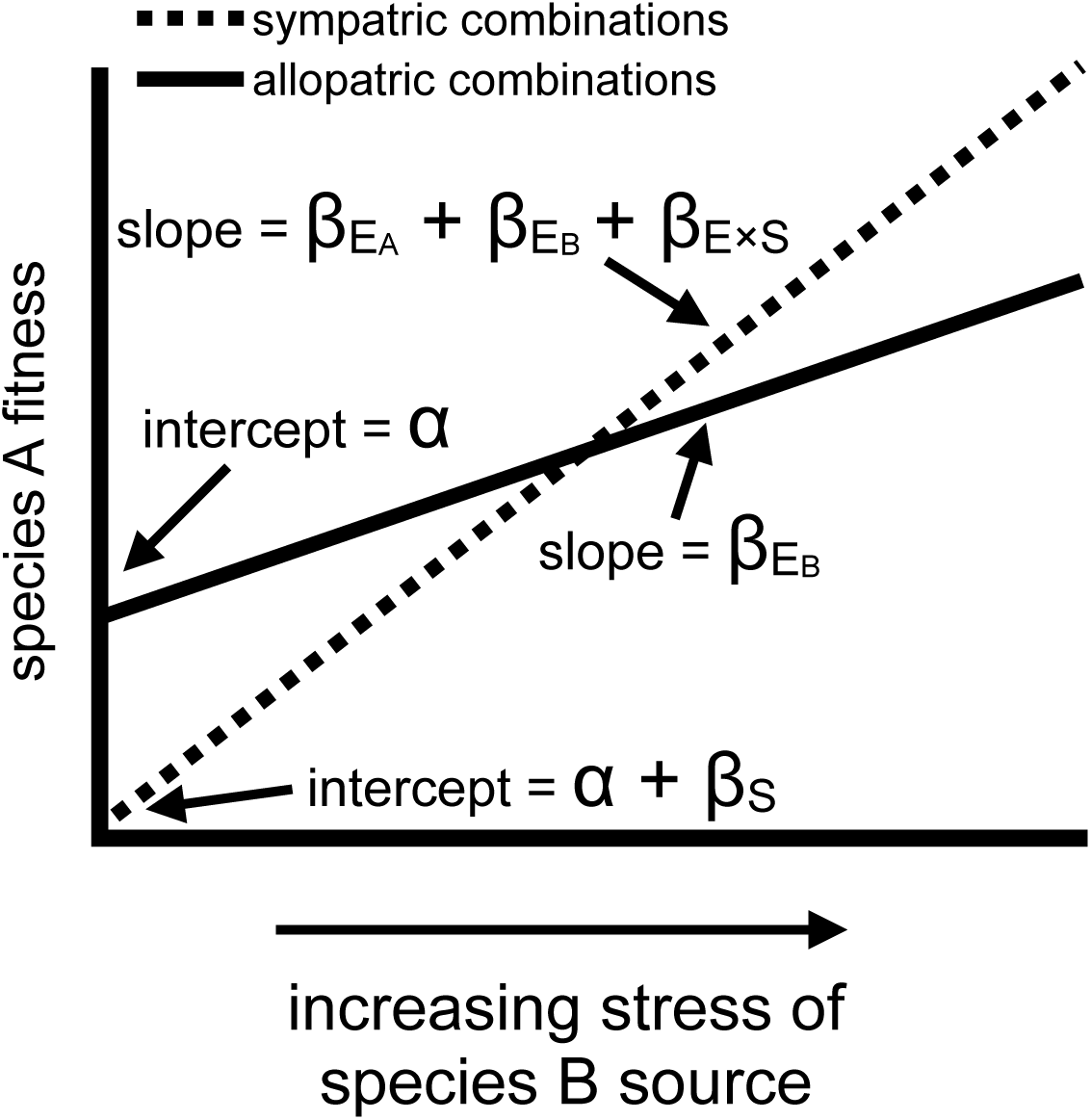
Here we relate model parameters to linear relationships between fitness and partner source, drawn here for the scenario in Figure 2 D. Generalized coevolutionary benefits are tested by the parameter *β_E_B__*, the slope of the allopatric comparisons (solid line). Specific coevolutionary benefits are tested by the parameter *β_E×S_*, which, when added to *β_E_B__* and *β_E_A__*, is the slope of the sympatric (dashed) line. β_S_ only affects the intercept of the sympatric line. *β_E×S_* alone describes the increasingly positive difference between allopatric and sympatric combinations as the source site becomes more stressful.

The focal partner source environment (here, the environment *E_A_*) is included to account for any main effects of population fitness along the gradient, as selection to reduce the fitness-limiting stress may not act only on interactions. For example, selection may increase tolerance of stress without interactions (Espeland and Rice, 2007), or low resource environments might select for smaller individuals than high resource environments. If such effects are large, they can cause over- or under-estimation sympatric effects (Blanquart et al., 2013). Since the slope of species A fitness along increasing source site stress of B partners is built from the sum of *β_E_A__*, *β_E_B__*, and *β_E×S_* (Figure 4), failure to account for *β_E_A__* can affect estimates of *β_E×S_* if fitness of Species A is positively or negatively correlated with the stress gradient. Estimating *β_E_A__* allows us to account for either of these other sources of correlation (see Blanquart et al., 2013).

Our above model assumes, and our figures (1,2,4) depict, a linear relationship between fitness and the environmental gradient. To assess whether non-linear effects of gradients are better descriptors of the effects on fitness of species interactions along gradients (e.g. Malkinson and Tielborger, 2010; Holmgren and Scheffer, 2010), and subsequent adaptation patterns, models with quadratic terms for *E_B_* and *E × S* should be compared with models using linear terms. Various types of non-linearity may be relevant, such as threshold or parabolic models, especially for interactions where peak mutualism or facilitation might be at mid-stress (see “Other considerations” below). Additional random effects that might be required, depending on the design, could include: family effects, block effects, or year effects (represented here as a generic *Z*, with parameter *β_Z_*).

### Interpretation of results

The predictions of CoCoA would be supported by the following outcomes: 1) if partners from more stressful sites provide greater benefits across all focal species populations than partners from less limiting sites (generalized benefits, *β_E_B__* significantly positive) and 2) if benefits that are provided to sympatric partners over allopatric partners are greater for populations from stressful sites (greater specific benefits, indicated by a significant and positive *β_E×S_*). When both *β_E_B__* and *β_E×S_* are significant and positive, both predictions of CoCoA are supported, and both allopatric and sympatric lines will have a positive slope (see Figure 4), but the sympatric line will be steeper (illustrated in Figure 2 D). For interactions that grade into facilitative commensalism (not depicted in Figures 2 and 4), we still expect to see increasing generalized and/or specific benefits, as for interactions that grade into mutualisms. However, such patterns should only be detected for the facilitated species.

Extensive gene flow between populations at stressful sites could result in more mutualistic partners from highly limited sites without increased local adaptation. For example, populations might experience isolation by environment more than isolation by distance (e.g. Sexton et al., 2016). This scenario would be indicated by the case that *β_E×S_* is non-significant and *β_E_B__* is significant and positive. The slope of the allopatric and sympatric lines would be identical (Figure 2B), or differences would be due only to patterns in fitness of the focal species across the gradient, *β_E_A__*, unrelated to species interactions.

This section has focused on the stressful ends of gradients and interactions that at least grade into mutualistic (+,+) or commensal (+,0) outcomes. A similar experimental design and model would be required for tests of CoCoA in antagonisms or at benign ends of the gradients. *β_S_* tests the main effect of sympatry, and is the intercept adjustment of the sympatric line relative to the non-sympatric line (Figure 4). This term reflects the difference between allopatric and sympatric pairings of A and B from benign sites. When this parameter is negative (as in Figure 2, B and D), it would indicate antagonistic adaptation in the non-focal species in benign sites. However, an estimate of *β_S_* that is positive or not different from 0 does not necessarily indicate a lack of antagonistic adaptation or co-adaptation, as adaptation in antagonistic interactions can generate non-significant effects (due to e.g. temporal or spatial variation in adaptation cycles, see below). Experimental evolution, especially with the ability to archive and resuscitate genotypes (in species with resting propagules), would allow detection of whether local mutualistic adaptation proceeds reciprocally (co-adaptation) or if one species alone produces all patterns of adaptation. CoCoA expects the same patterns in increasing specific benefits with stress regardless of whether responses to selection are reciprocal (local co-adaptation) or restricted to one species (adaptation only); benefits measured in sympatric pairs do not separate contributions of adaptation in each species.

### Other considerations

A non-trivial matter is how the gradient is defined and identified. Specifically, for CoCoA to hold, not only must sites be stressful, but interactions between partners must ameliorate the stress. CoCoA will be predictive when conditions for the SGH and limiting resource conditionality are met: when a stress ranges from non-limiting to strongly limiting of fitness and is ameliorated by interaction between the focal species (He and Bertness, 2014). CoCoA will further be most predictive when gene flow is sufficiently restricted to allow local adaptation and there is genetic variation on which selection can act in both partners. CoCoA will be less informative across weak, non-limiting, or multiple co-occurring gradients, where importance of interactions to fitness is less predictable (He and Bertness, 2014).

While extensive research on the SGH in plant-plant interactions generally supports the prediction of increasing facilitation with stress (He et al., 2013), peak facilitation may occur in sites with moderately, rather than extremely limiting stress (Michalet et al., 2006; Holmgren and Scheffer, 2010; Malkinson and Tielborger, 2010). Such intermediate peaks could be generated by nonlinear relationships between benefits (or costs) and abiotic gradients (Holmgren and Scheffer, 2010), or by low density of individuals at high stress sites causing missed interactions (Travis et al., 2006). Intermediate peaks appear to fit best in interactions that grade from increasing to decreasing access to a shared limiting resource (Maestre et al., 2009; Michalet et al., 2014), as opposed to interactions with differing limiting resources between partners. Plant-pollinator benefits also can show intermediate peaks, because relationship between pollination limitation and fitness limitation often changes non-linearly across environments (Haig and Westoby, 1988; Maron et al., 2014). Non-linearity also makes sense in light of the fact that there may be little interactions can do in the face of extreme stress, and if they no longer ameliorate the stress, then selection will no longer favor investment in the interaction. Peaks for positive outcomes in moderately stressful conditions, regardless of mechanism, have the consequence for CoCoA that mutualistic adaptation and co-adaptation would also peak at moderately stressful conditions, in which case, non-linear models for fitness across stress gradients would be needed (see “A linear model framework” above).

Source site differences along the stress gradient may affect fitnesses of partner pairings. In studies of climate adaptation, functions of environmental distance transfer from source site to experimental site better predict success than experimental site environment alone (Wang et al., 2010). If species interactions have analogous dynamics, instead of or in addition to CoCoA, then such transfer functions between source sites of experimental combinations would determine their ability to mutually benefit from each other, rather than dynamics of local and mutualistic adaptation. For example, we combine CoCoA effects with the transfer function of Wang et al. (2010), by adding squared source environment terms and an interactive slope, *β_E_A__*×*E_B_*:

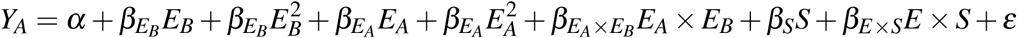

The quadratic model of Wang et al. (2010) here now implies that there is either an optimal environmental distance (i.e. potentially 0 for mutualisms), or least optimal distance between sources of partner populations along the environmental gradients. As the distance between population pairs increases, fitness effects on the focal species either increase or decrease, depending on parabola sign. As before, the addition of sympatry, and sympatry-by-environment effects just add the local (co)-adaptation effects we have discussed here, and generalized benefits are captured by the linear source environment terms. Power to estimate such transfer functions, would be improved by many more population comparisons than demonstrated in Figure 3 (see Wang et al., 2010).

We have focused our tests and predictions around conditions that predict mutualistic coevolution (high stress) because coevolutionary patterns from antagonisms (predicted in benign conditions) are more difficult to detect. Character displacement in -,- interactions is notoriously difficult to document (Stuart and Losos, 2013). Similarly, in +,- interactions, one species may be “winning” the battle and appear locally adapted at a single timepoint, but the winning species is likely to vary across both timepoints and space as evolution in the other species counteracts “gains” (e.g. Van Valen, 1974; Gandon and Michalakis, 2002; Nuismer, 2006). Running this experiment multiple times from populations collected at different timepoints (see Decaestecker et al., 2007), or across experimental evolution (see Pascua et al., 2011) would allow differentiation between drifting variation in sympatric effects and Red Queen dynamics. Long-term sampling of trait changes and genotypes (Dybdahl and Lively, 1998; Decaestecker et al., 2007), as well as long term partner removal experiments (Stuart and Losos, 2013), have also proven to be effective tools for detecting antagonistic coevolution, and would be equally useful for testing CoCoA predictions in antagonistic interactions. Regardless of the test, conclusions must be based on degree of trait change or rate of evolutionary dynamics across both abiotic gradients and time.

## Existing literature pertinent to CoCoA

In reviewing the literature, we found a number of studies in which most, but not all, of the criteria required to evaluate CoCoA have been tested, but only one study that has addressed all criteria. These studies, however, have some evidence related to the predictions of CoCoA.

Experimentation on plant-microbe interactions offer the most complete tests. The outcomes of interactions between plants and rhizosphere biota (a diverse community of microbes living in and near roots Hiltner (1904)) are highly influenced by environments (e.g. Zhu et al., 2009; Smith and Read, 2008; Lau and Lennon, 2012). Limiting soil nutrients have frequently been identified as the potential driver of the evolution of interactions with soil rhizosphere microbes (Johnson, 1993; Schwartz and Hoeksema, 1998; Kiers and van der Heijden, 2006; Bever, 2015), and metaanalysis finds local adaptation in plants and mycorrhizal fungi to be common but not universal (Rúa et al., 2016).

Johnson et al. (2010), which met all of the above criteria, found mutualistic local adaptation between a grass and its associated arbuscular mycorrhizal fungi across a phosphorus gradient. Plants are generally known to derive increased benefits from interacting with these fungi in low phosphorus conditions (Smith and Read, 2008). Fungi sourced from low phosphorous sites were more beneficial across plants but provided even greater benefits to sympatric plants (Johnson et al., 2010), supporting both the specialized and generalized benefits predictions of CoCoA. However, as only three sites were sampled, we remain cautious of inferring strong support for CoCoA. Other studies sample outcomes along environmental stress gradients, but do not explicitly include sympatric and allopatric partners to evaluate the nature of benefits or local adaptation (specific or generalized). Barrett et al. (2012) cross-inoculated acacias and microbes sampled along a soil nitrogen gradient (likely a limiting stress gradient), and found that the effects of soil microbes sampled from low nitrogen sites provided the greatest benefit to acacias. In a study of plants and nitrogen-fixing bacteria, bacterial genotypes sampled from high nitrogen sites (in which nitrogen is less limiting to plants) similarly provided less benefits than genotypes from low nitrogen sites (Weese et al., 2015).

In many ant-plant mutualisms, ants protect plants from herbivory and receive food from the plant. In drier sites, plants are limited by both water and herbivory costs, and ants are likely limited by plant-fixed carbon (Pringle et al., 2013). In such dry, limited sites, ants invest more in plant defense, reducing herbivory limitation, and plants allocate more carbon to ants, increasing ant colony size (Pringle et al., 2013). In Pringle et al. (2013), lower water sites were limiting for a plant host because insufficient water increased the risk of plant death from herbivory. This example documents both the limiting gradient, which is ameliorated by the interaction for both partners, and greater reciprocal mutualistic benefits at the stressful portion of the gradient. It remains to be seen whether these benefits are adaptive differences or plastic behaviors, and whether they are generalized or specific.

Plant-plant interactions across mesic-arid gradients range in outcome from antagonistic to facilitative as aridity increases (He et al., 2013), leading to the prediction of CoCoA that adaptation to competitors would be greatest in mesic sites and adaptation of beneficiaries to facilitators greatest in arid sites. Existing evidence does not reject CoCoA, but also does not offer complete tests: genotypes of plants from mesic (benign) sources were least affected by competition in multiple systems (Liancourt and Tielborger, 2009; Liancourt et al., 2013), and another study found greater evidence for plant local adaptation in mesic sites when neighbors are included (Ariza and Tielborger, 2011). However, a test of adaptation in plant-plant interactions from a different stress gradient (soil chemical stress) suggests that adaptive increases in stress-tolerance may be more important than adaptive increase in benefit from facilitation (Espeland and Rice, 2007).

Bacteria-phage systems at the conditions least stressful for bacteria (high nutrients) show strongest local adaptation (receipt of specific benefits) of phages to host bacteria (Pascua et al., 2011). Pascua et al. (2011) also showed increasing overall infectivity and resistance in high nutrient conditions, suggesting greater trait escalation. Another study found the reciprocal expectation: less stressful conditions for bacteria led to evolution of increased defense traits in bacteria when phages and bacteria were permitted to evolve (Zhang and Buckling, 2016).

Increased trait escalation at high productivity (indicating low plant stress) has also been found in camellia-weevil antagonisms, where camellia defensive and weevil offensive (Toju et al., 2011) traits appear to have escalated more. However, it is clear that not all plant-herbivore interactions change in outcome the same way across plant productivity gradients (Maron et al., 2014). In sites where prey are physiologically less limited, defensive traits appear to have escalated more in newt-predator (Stokes et al., 2015) and squirrel-rattlesnake (Holding et al., 2016) predator-prey antagonisms.These systems show some of the patterns CoCoA would predict, but whether stress-gradients led to these patterns, or whether patterns reflect adaptation to interactions must still be tested.

In contrast, we found only one study with evidence in conflict with CoCoA predictions. Across a gradient of increasingly cold conditions, plants show no local adaptation with rhizosphere biota and no evidence of increasing benefits from colder sourced biota (Kardol et al., 2014). While the extreme cold is very likely to be stressful, and the ability of interactions plant-rhizosphere biota to reciprocally ameliorate effects of extreme cold also likely (Zhu et al., 2009), they were not tested.

In sum, while current evidence offers mixed support, only very few tests of CoCoA exist. Complete tests of CoCoA are within reach in many more systems, and evidence above suggests that complete tests of CoCoA as outlined above would be worthwhile.

## Discussion

Economic and stress-gradient models of conditionality in species interactions predict shifts in species interactions from more negative outcomes to more positive outcomes as environmental stress (Bertness and Callaway, 1994; Brooker and Callaghan, 1998; Malkinson and Tielborger, 2010) or resource limitation (Johnson, 1993; Schwartz and Hoeksema, 1998) increases (see Figure 1).

We present here an extended hypothesis for the evolutionary consequences of these models of ecological conditionality, which we term Co-adaptation to Conditionality across Abiotic gradients (CoCoA). CoCoA predicts increasingly strong (co-)evolutionary dynamics where conditionality models predict increasingly positive or increasingly negative interaction outcomes. At stressful sites where partners mutually enhance each others’ fitness, or one partner receives increasing benefits, mutualistic co-adaptation and adaptation are predicted to dominate, respectively. In benign sites where the interaction shifts towards parasitic, or mutually negative, CoCoA predicts intensification of evolutionary dynamics: Red Queen (or similar) coevolutionary scenarios for parasitic outcomes, and adaptation to avoid interactions such as character displacement (Pfennig and Pfennig, 2009) or habitat partitioning (Martin, 1998), for mutually negative outcomes. Between these extremes, in intermediate stress or benign environments, interaction outcomes approach neutrality, leading to predictable zones of no adaptation.

Other models of co-adaptation (Johnson, 1993; Schwartz and Hoeksema, 1998; Thrall et al., 2007; Bever, 2015) and behavioral models (Revillini et al., 2016), focus on environmental gradients. Like CoCoA, some models (Johnson, 1993; Schwartz and Hoeksema, 1998; Bever, 2015) predict that selection in resource-limiting environments should favor increased benefits provided to partners in the mutualism. Alternatively, Thrall et al. (2007) make predictions based on levels of environmental productivity and biological diversity. CoCoA differs from these models in its focus on adaptation patterns in both partners, its inclusion of fitness-limiting stresses beyond resources, and thus its applicability to a wide variety of conditional interactions.

CoCoA implies that selection for specialization may be common at both ends of the stress gradient continuum, i.e. in both antagonistic and mutualistic interactions. While it is generally accepted that parasitism often promotes specialization and increases the rate of evolution (Paterson et al., 2010), it is debated whether mutualism commonly imposes selection for specialization (Thompson, 2005). There is, however, some evidence that mutualism can be at least as strong a driver for specialization as parasitism (Kawakita et al., 2010), and mutualists may evolve at equal or faster rates than non-mutualist sister lineages (Rubin and Moreau, 2016).

## Concluding Remarks

As climatic conditions become more extreme and stressful under global change (Pachauri et al., 2014), we predict that adaptation to changing environments may be heavily influenced by biotic interactions. Numerous studies have focused on single species processes that limit ranges, such as source-sink dynamics or maladaptive gene flow (see Sexton et al., 2009, for review), but our CoCoA hypothesis suggests more research on multi-species dynamics may be fruitful (Sexton et al., 2009; van der Putten et al., 2010).

CoCoA contributes to a growing body of literature highlighting the importance of biotic interactions in determining limits of species distributions across abiotic gradients (e.g. HilleRisLambers et al., 2013; Afkhami et al., 2014), even in climatically stressful environments (e.g. Brown and Vellend, 2014) where abiotic variables have often been thought to be of greater importance (Brown et al., 1996; Hargreaves et al., 2014). Biotic filters on abiotic variables that exacerbate or ameliorate abiotic effects may thus have widespread consequences for range shifts and other responses to global change.

## Acknowledgements

AO was supported by NSF GRFP grant DGE-1148897 and NSF grant DEB-0919559 to SYS. J.R.I. would like to thank support from NSF Plant Genome (project I0S-1238014) and USDA (Hatch project CA-D-PLS-2066-H). R.J.H.S would like to thank support from CONACYT (CB15-25401). The authors would like to thank the Coop, Schmitt, Strauss, and Ross-Ibarra labs at UC Davis for discussion.

